# Individual level reliability of PAS-induced neural plasticity

**DOI:** 10.1101/130856

**Authors:** Yeun Kim, Jacqueline P. Ngo, Choi Deblieck, Dylan J. Edwards, Bruce Dobkin, Allan D. Wu, Marco Iacoboni

**Author notes:** Email addresses (Yeun Kim), (Jacqueline P. Ngo), (Choi Deblieck), (Dylan J. Edwards), (Bruce Dobkin), (Allan D. Wu), (Marco Iacoboni).

## Abstract

**Objective:** We assessed the individual level reliability of neural plasticity changes induced by paired associative stimulation (PAS), which combines peripheral nerve stimulation with transcranial magnetic stimulation to induce short-term plastic changes in the brain.

**Methods:** For 5 consecutive weeks, motor evoked potentials (MEPs) of 8 healthy subjects were acquired every 10 minutes post-PAS intervention for a period of 60 minutes. The post-PAS MEPs were evaluated against base-line MEPs using permutation and Kolmogorov-Smirnov tests to determine whether the MEP magnitudes changed after PAS. Moreover, various sample sizes of the MEP data were used to deduce the minimum number of MEPs needed to reliably detect individual propensity to neural plasticity.

**Results:** Group analysis exhibited significant increase in post-PAS MEPs, confirming previous results. While high between-sessions variability was observed at individual level, data show that between 40 to 50 MEPs can reliably assess each subject’s responsiveness to PAS. Subjects exhibited three different plasticity patterns: in the modulated hemisphere only, both hemispheres, or neither hemisphere.

**Conclusions:** PAS can reliably assess individual differences in neural plasticity.

**Significance:** A marker of individual plasticity may be useful to predict the effects of a motor rehabilitation, drug or other intervention to increase recovery of function after brain injury.

**Highlights:**

- Paired associative stimulation (PAS) assesses neural plasticity non invasively.
- The study shows how PAS can reliably determine individual differences in plasticity.
- PAS may be used to predict intervention outcome or individualize treatment dose.

## 1. Introduction

Neural plasticity refers to the brain’s lifelong ability to adapt to change by altering its functional organization, via processes including neuronal growth, neurotransmitter modulation, and alteration of synaptic connections (Nudo, 2006). It occurs in response to subject′s activity, such as exercise or skill acquisition, and other kinds of events, as for instance brain injury, or the administration of pharmaceutical drugs (Cotman and Berchtold, 2002; Nudo, 2003; Kleim et al., 2003; McEwen and Chattarji, 2004; Nitsche et al., 2012). Neuromodulation with non-invasive stimulation techniques, represents a contemporary approach to assess brain plasticity responsiveness in the intact or damaged human brain. One such paradigm is paired associative stimulation (PAS), a non-invasive method of brain stimulation, in which an electric stimulus is applied to a peripheral nerve (PNS) before transcranial magnetic stimulation (TMS) is delivered to the contralateral primary motor cortex (M1). PAS can increase or decrease corticospinal excitability (CSE) with a mechanism thought to be similar to spike timing-dependent plasticity, in which the order and temporal interval of neuronal firing adjusts the strength of connections between neurons. An inter-stimulus interval (ISI) of 25 ms between PNS and TMS is typically linked with increased CSE, while shorter ISIs as for instance 10 ms are typically associated with decreased CSE (Carson and Kennedy, 2013). CSE serves as a marker of long-term potentiation or long-term depression-like plasticity, which can be induced by PAS and assessed by changes in motor evoked potential (MEP) amplitude post intervention relative to baseline (Stefan et al., 2002).

Thus, by allowing us to non-invasively probe cortical plasticity in human subjects, PAS serves as a potentially useful tool to explore how the corticospinal system might be responsive to conventional interventions such as motor learning through practice of skills, or pharmacologic manipulation that aim to augment learning, plasticity and behavior, especially after stroke or injury to the motor pathways.

Previous PAS studies targeting LTP-like effects have shown that group data reveal a reliable increase in CSE (Wischnewski and Schutter, 2016; Nitsche et al., 2007; Rajji et al., 2011). However, reproducibility at the single subject level is limited. Neurologically healthy individuals display wide variability in their response to PAS and other types of brain stimulation, as assessed by retesting (Fratello et al., 2006). Multiple factors contribute to this variability. Within-subject factors include priming, prior and ongoing voluntary motor activity, aerobic exercise, attentional state during intervention (Stefan et al., 2004), time of day (Sale et al., 2007), and intrinsic neuronal activity (Nitsche et al., 2007). Between-subject factors include age (Sale et al., 2007; Müller-Dahlhaus et al., 2008), gender (Tecchio et al., 2008), cortical thickness (Conde et al., 2012; List et al., 2013), and genetic polymorphisms (Kleim et al., 2006; Cheeran et al., 2008; Missitzi et al., 2011). Finally, multiple drugs are reported to alter plasticity responses, such as tianeptine (McEwen and Chattarji, 2004), ketamine (Wang et al., 2015) and fluoxetine (Chollet et al., 2014).

A method of assuring reliable individual responses to the PAS intervention would be highly valuable. Indeed, reliable individual responses to PAS would allow to test the efficacy of drug, practice, noninvasive brain stimulation, cellular and other treatments to enhance motor recovery in individual patients with upper extremity neurological impairment. In this paper, we hypothesize that subject-level reliability of PAS is achievable with a sufficient number of observations, and propose a methodological recommendation for future PAS studies that aim at assessing individual differences in plasticity.

## 2. Methods

### 2.1. Participants

Ten healthy participants (two males, eight females, aged 19-36 years, M=24.4, SD=6.15) were recruited for the study. Each subject participated in five stimulation sessions, spaced one week apart. Two subjects were excluded from final data analysis due to technical issues leading to missing baseline motor evoked potential (MEPs) data from two different sessions. Data analysis was performed on the remaining eight subjects (two males, six females, M=24.25, SD=6.18). The study was approved by the UCLA Institutional Review Board, and written informed consent was obtained from all participants.

### 2.2. Electromyography recording

All participants were seated comfortably in an armchair. Surface electromyography (EMG) electrodes (Delsys 2.1, Delsys, Inc) were attached to the first dorsal interosseus (FDI) muscle of each hand to measure MEPs. The electrical activity of the muscles was amplified and bandwidth filtered between 20 and 450 Hz (Bagnoli EMG, Delsys, Inc), digitized with a sample rate of 5 kHz (CED 1401 running Signal V4 software, Cambridge Electronic Design, Cambridge, UK) and stored for off-line analysis using custom Matlab routines.

### 2.3. Transcranial magnetic stimulation

TMS was delivered through a figure-eight shaped coil (70mm diameter each coil) connected to a Magstim 2002 magnetic stimulator (Magstim, Whitland, Dyfed, UK). TMS pulses for PAS were always delivered over the left motor cortex in each subject. However, pre and post PAS effects (corticospinal excitability, CSE) were assessed by TMS pulses applied over both left and right primary motor cortices in succession. For each CSE assessment, TMS pulses were applied to left motor cortex first for odd-numbered subjects, while TMS pulses were applied to the right motor cortex first for even-numbered subjects.

The coil was held with the handle pointing posteriorly at a 45-degree angle down the sagittal plane and moved systematically across the scalp to identify the optimal position and orientation of the coil for the “hot spot” for each motor cortex. The “hot spot” was defined as the point at which TMS consistently produced MEPs of maximum amplitude from the contralateral first dorsal interosseus (FDI). At the beginning of each session, the “hot spot” was identified, recorded onto a stereotactic system (Brainsight, Rogue Research, Montreal, Canada), and marked with a pen on the subject’s scalp. The mark on the scalp helped initial repositioning across time points within a session and visual feedback from Brainsight ensured that coil positioning was kept at optimal location during stimulation. For this hot spot, the resting motor threshold (rMT) was established, defined as the lowest stimulus intensity required to evoke an MEP of 0.05 mV or higher in at least 3 out of 6 consecutive trials in the relaxed FDI.

The TMS intensity consistently producing MEPs in the 0.5 to 0.75 mV range in the relaxed FDI was subsequently determined for each motor cortex and was used as the test intensity to measure baseline corticospinal excitability (CSE). Previous PAS studies have typically used 1 mV as the target amplitude to determine TMS intensity. Since this individual level reliability study was designed having in mind its applicability to studies in clinical populations, a lower amplitude was deemed more feasible for these future studies. Note that this lower amplitude does not affect PAS itself (whose intensity is based on rMT) and cannot possibly bias the main outcome measure of this study, which is individual level reliability of PAS response.

To obtain one block of 10 baseline MEPs, TMS pulses were given at random interstimulus intervals (ISI) between 8-12 seconds. This procedure measured baseline corticospinal excitability. To control for attention, subjects were asked to look at the target muscle, count the number of times they felt their muscle twitch, and report this number at the end of the protocol (Stefan et al., 2004). Data acquisition followed a method which routinely excluded the first two MEP measures (O’shea et al., 2007) and kept the next 10 MEPs that passed these three criteria: (1) minimum peak-to-peak amplitude of 0.5 mV (this was the lower value of the range of amplitudes used to determine TMS intensity subsequently used for baseline MEPs); (2) latency of no less than 20 ms from stimulation; (3) no presence of voluntary movement during the pre-stimulus phase. The EMG signal was monitored visually throughout the entire course of stimulation and reviewed offline during data analysis. The hot spot location, rMT, and baseline MEP assessment were repeated in the contralateral hemisphere before the PAS intervention.

### 2.4. Paired associative stimulation

Paired associative stimulation (PAS) consisted of repetitive single electric stimulation (square wave, 0.2 ms duration) to the right ulnar nerve delivered using a DS3 Isolated Current Stimulator (Digitimer, UK) followed by single TMS pulse over the left hemisphere hot spot with an interstimulus interval (ISI) of 25 ms (Player et al., 2012). 200 pairs of stimulation were applied at 0.25 Hz. The ulnar nerve was stimulated at an intensity of 300% perceptual threshold (Stefan et al., 2000) and TMS intensity was set at 130% rMT (Player et al., 2012).

### 2.5. Post PAS assessment

To obtain each post-PAS block of MEPs, TMS pulses were delivered with the same intensity used for baseline MEP assessment to each hemisphere immediately after PAS and in 10-minute intervals for a period of 60 minutes. At each time point, blocks of 10 MEPs with a random ISI of 8-12 seconds were acquired. To control for attention, subjects were asked again to look at the target muscle, count the number of times they felt their muscle twitch, and report this number at the end of the protocol (Stefan et al., 2004). Data acquisition was as for baseline MEPs (see 2.3). The study protocol is illustrated in figure 1.

**Figure 1:**
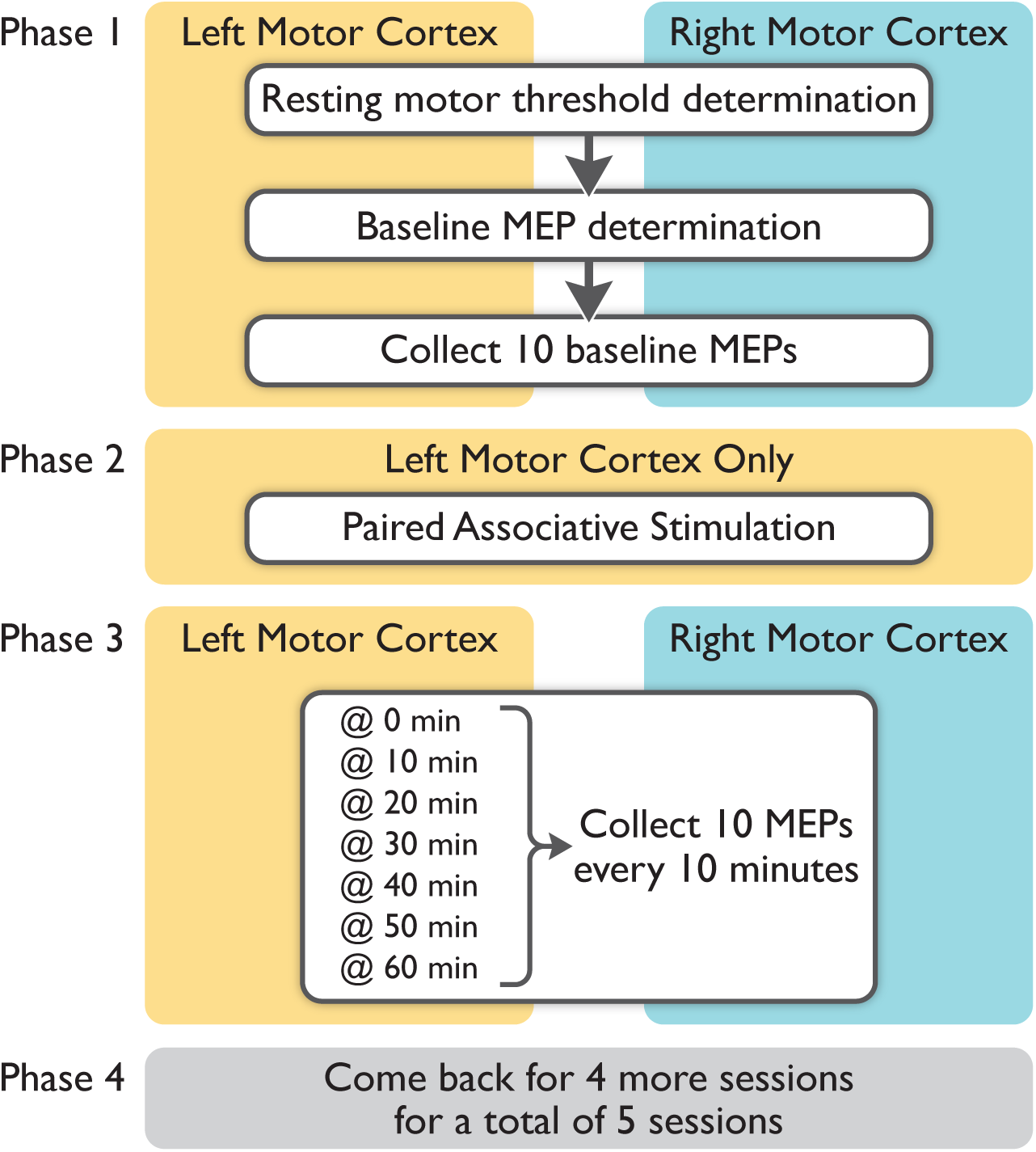
Schematic diagram of study protocol.

### 2.6. Data Processing

The data was first corrected for direct current offset to ensure that the EMG signal within the pre-stimulus phase had an average of 0 mV. The window for the calculation of the MEP waveform began at 20 ms after the stimulus onset and had a duration of 50 ms. Then, the area under the waveform was computed by taking the integral using the trapezoidal rule after the data was rectified. The digital signals were not further smoothed or filtered to preserve the integrity of the waveforms.

We also calculated the local minimum and maximum of the waveforms to compute the peak-to-peak amplitude. However, we chose to use area-under-the-curve (AUC) measures, because they were able to better characterize the overall magnitude of multi-modal waveforms (Luck, 2005).

### 2.7. Statistical analysis

#### 2.7.1 MEP inclusion for data analysis

We compared three different methods of MEP inclusion for subsequent data analysis. We did so to determine whether different inclusion criteria might potentially bias final results.

1. Ten MEP measures after excluding the first two MEP measures at each time point. These measures must also satisfy all the following criteria (verified visually during data acquisition and confirmed at data processing stage):

- Latency period of 20ms after stimulation
- Minimum peak-to-peak amplitude of 0.5 mV
- No voluntary activity within the pre-stimulus phase
2. The first ten MEPs at each time point.
3. All MEPs at each time point.

To compare these three methods of MEP inclusion in subsequent analyses, we calculated the area under the curve for each MEP, then took the median value at each time point. Since the Kolmogorov-Smirnov (KS) test showed a non-normal distribution, we log-transformed the data and performed a twoway repeated measures ANOVA test. We observed no significant difference between the MEP values across the three methods. We took this result as suggesting that choosing one method over another would not significantly bias final results. Therefore, we adopted method (1) for data analysis, which is commonly used in PAS studies (Müller et al., 2007). This method was also the one that aligned with our data acquisition protocol.

#### 2.7.2. Group level analysis

As mentioned previously, the ability of PAS to induce changes in cortical excitability has been well-reported in previous studies using group data (Stefan et al., 2000; Wolters et al., 2003; Ziemann et al., 2004; Morgante et al., 2006; Weise et al., 2006). Thus, we performed a repeated measures ANOVA on our group data to verify that it replicated the previously reported findings.

Each subject’s median baseline MEP was used as the normalization factor for their post-PAS MEPs. No normal distribution was found when the KS test was applied, so Box-Cox was used to normalize the data (*λ*=-0.0606 for LM1-RFDI; *λ*=0.1414 for RM1-LFDI). We chose to use a power transform instead of a log transform because the power transform method was able to generate a distribution that better fits normality according to both the Shapiro-Wilk and Anderson-Darling test. Then, a two-way repeated measures ANOVA was performed to test for significant time point, session, and time point by session interaction effects.

Next, to determine whether there was a (1) significant linear trend of MEPs change and (2) if this was different between hemispheres, we performed contrast analysis using orthogonal polynomial contrasts in both hemispheres, as well as tested for the interactive effect between time points and hemispheres.

#### 2.7.3. Individual level analysis

The individual data was normalized and power transformed using the same method as the group level analysis. ANOVA was applied to test for time point by session interaction effects within individual subjects. For each subject, the tests were separated into two families: one for tests performed on the modulated side, and one for tests performed on the non-modulated side. False discovery rate (FDR) was applied to each family of tests to correct for family-wise error rate (FWER).

Since these individual ANOVAs demonstrated time point by session interaction at the subject level (see 3.2.1), we performed the following tests to investigate whether each individual session had an insufficient number of data points to demonstrate a reliable change at the individual subject level.

We repeated the same non-parametric statistical tests while varying the number of MEPs analyzed for each time point for each subject by combining random combinations of the 10 normalized MEPs from a set of sessions between 1 and 5, each of which consisted of 10 normalized MEPs. For instance, we first performed tests using 10 MEPs from one session. Next, we applied the same test using 20 MEPs (combined from two random sessions); we repeated this step two more times using different sets of 2 sessions. Then, we continued to perform the tests using the same method with 30, 40, and 50 MEPs to determine whether we could observe a consistently reliable significant difference in post-PAS MEPs with larger samples of MEP data. Kolmogorov-Smirnov tests were applied to each time point to determine whether the MEP distribution were significantly different from the baseline MEPs. Normalized MEP values from all sessions were used for these analyses. FDR was applied to correct for multiple comparisons.

Moreover, we used permutation tests to simulate a normal distribution under the null hypothesis of no significant difference in MEP AUCs relative to baseline. We performed 5000 iterations, and considered a p-value *<*0.05 to be a measure for reliable improvement.

Lastly, we compared the power factor for sample sizes of 40 and 50 MEPs at each time point, since the KS and permutation tests demonstrated that these sample sizes were the most reliable at individual subject level (see 3.2.3). To obtain the effect size for the 40 MEP sample data, the effect sizes for all possible combinations of 40 MEPs (in groups of ten, by session) were calculated and averaged. Then, power was calculated for each sample size. Moreover, to determine whether there is a significant difference in power between 40 and 50 MEPs, a t-test and a non-parametric permutation test were applied at each time point across all subjects.

All statistical analyses were performed in R (v3.2.3) (https://www.r-project.org).

## 3. Results

### 3.1. Group level analysis

#### 3.1.1. Interaction effects between session and time point

The two-way repeated measures ANOVA showed no significant session by time point interaction effect (p=0.5696), suggesting that there were no reliable week-to-week changes in the temporal profile of the group response to PAS (Figure 2, Table 1). However, the data did exhibit a significant session effect (p=0.0013); the meaning of this finding is unclear and does seem rather orthogonal to the aim of this study.

**Figure 2:**
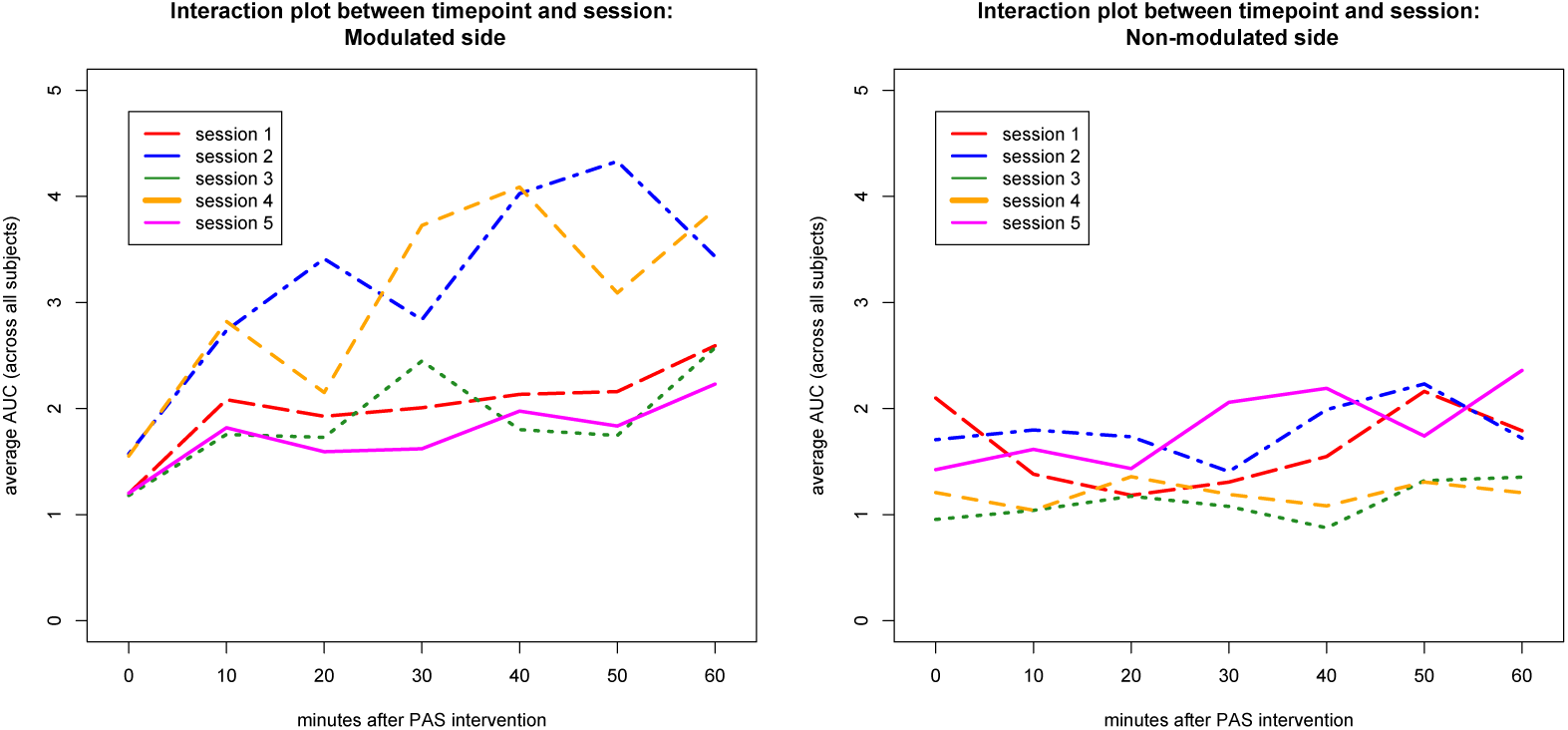
Interaction plots of LM1-RFDI (left) and RM1-LFDI (right). There was no significant session by time point interaction in either hemisphere.

**Table 1:** F-values and p-values from two-way repeated measures ANOVA performed on data from all subjects. Two ANOVAs were performed to test for the presence of interaction effects between session and time point as well as hemisphere and time point, while controlling for inter-subject variability and hemisphere.

| Predictor variables | F-value | P-value |
| --- | --- | --- |
| Session | 4.538 | 0.0013 |
| Time point | 0.134 | 0.7142 |
| Subject | 6.126 | $< 2 \times 10^{-16}$ |
| Hemisphere | 99.835 | $6.6 \times 10^{-07}$ |
| Session x Time point | 0.733 | 0.5696 |
| Hemisphere x Time point | 11.943 | 0.000591 |

#### 3.1.2. Time point and hemisphere effects

While the ANOVA revealed no significant time point effect (p=0.7142), it exhibited a significant hemisphere effect (p=6.6x10^-07^). These results suggest that the behavior of the MEPs differs between the modulated hemisphere and the non-modulated hemisphere. Thus, we performed an additional ANOVA and found significant interactive effects between time point and hemisphere (Table 1). To test for the presence of linear relationships within each hemisphere, we performed a contrast analysis.

#### 3.1.3. Linear trend

Contrast analysis exhibited a significant interaction between linear trend and hemispheres while controlling for sessions (p=0.000946, F=7.394). After testing for the presence of linear relationships in each hemisphere independently, the contrast analysis verified a highly significant linear trend (p=0.000103, t=-3.942) in the modulated (LM1-RFDI) side and a linear trend only approaching significance (p=0.0511, t=1.959) in the non-modulated side (RM1-LFDI) (see Figure 3). Importantly, since the lambda coefficient for LM1-RFDI’s power transformation was negative, the true, non-transformed sign of the contrast analysis’ slope is positive.

**Figure 3:**
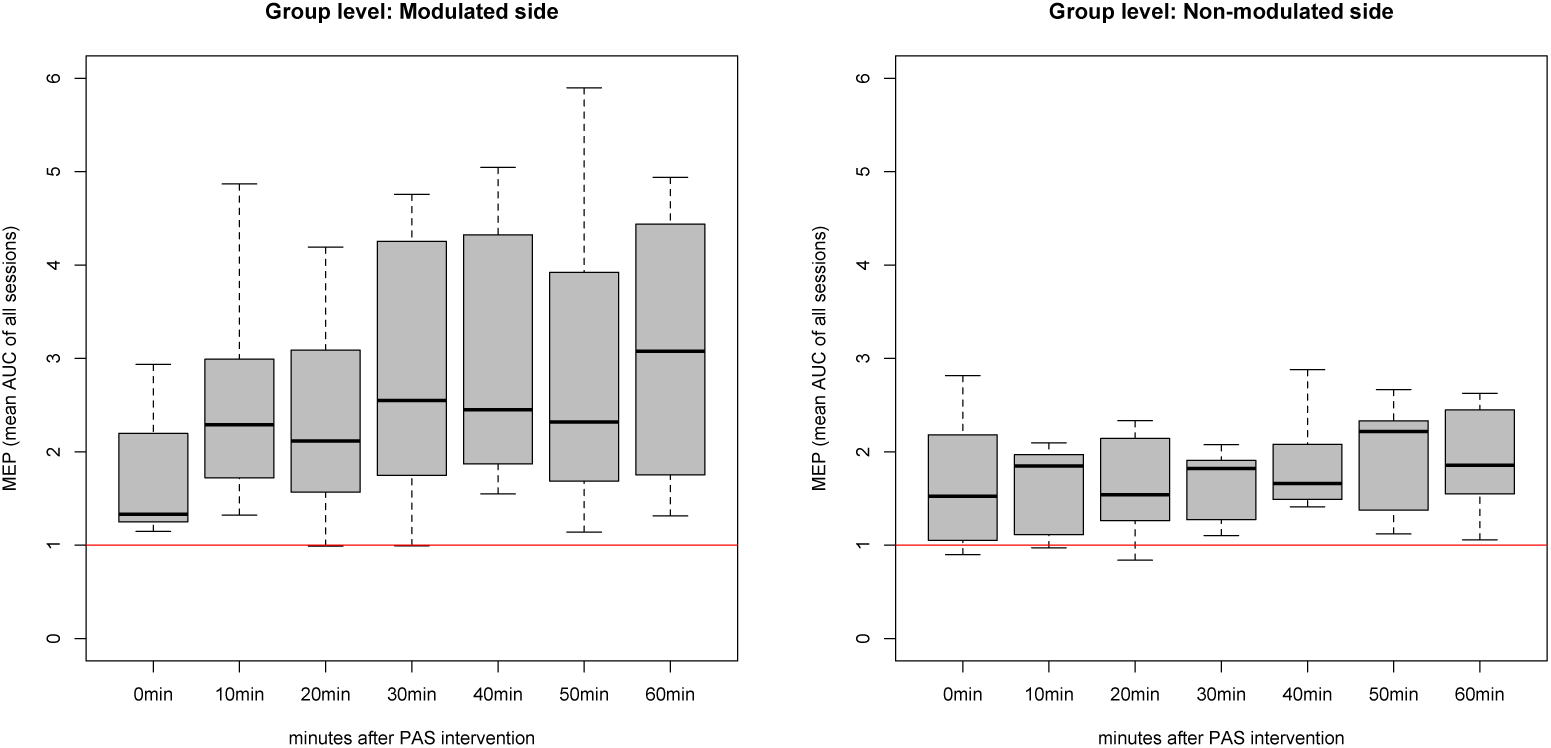
Each box plot contains all median MEP AUCs, normalized to each subject’s baseline, then averaged from all five sessions, for all subjects. Both groups exhibited significant session effects. Contrast analysis verified a positive linear trend (p=0.000103, t=-3.942) in the modulated side, while it only approached significance (p=0.0511, t=1.959) in the non-modulated side.

### 3.2. Individual level analysis

#### 3.2.1. Interaction effects between session and time point

At the intra-subject level, all subjects exhibited significant time point by session interaction effects in the modulated hemisphere (LM1-RFDI) (Figure 4). All subjects but pas03 (p=0.0799) exhibited significant time point by session interaction effects in the non-modulated hemisphere (see Figure 5). The fact that nearly all subjects show fairly similar interaction effects in both hemispheres suggests that data from individual sessions in individual subjects is rather noisy, perhaps due to an insufficient number of data points. This led us to the subsequent analyses that tested the minimal number of MEPs that reliably identify a response at the individual subject level and detect individual differences.

**Figure 4:**
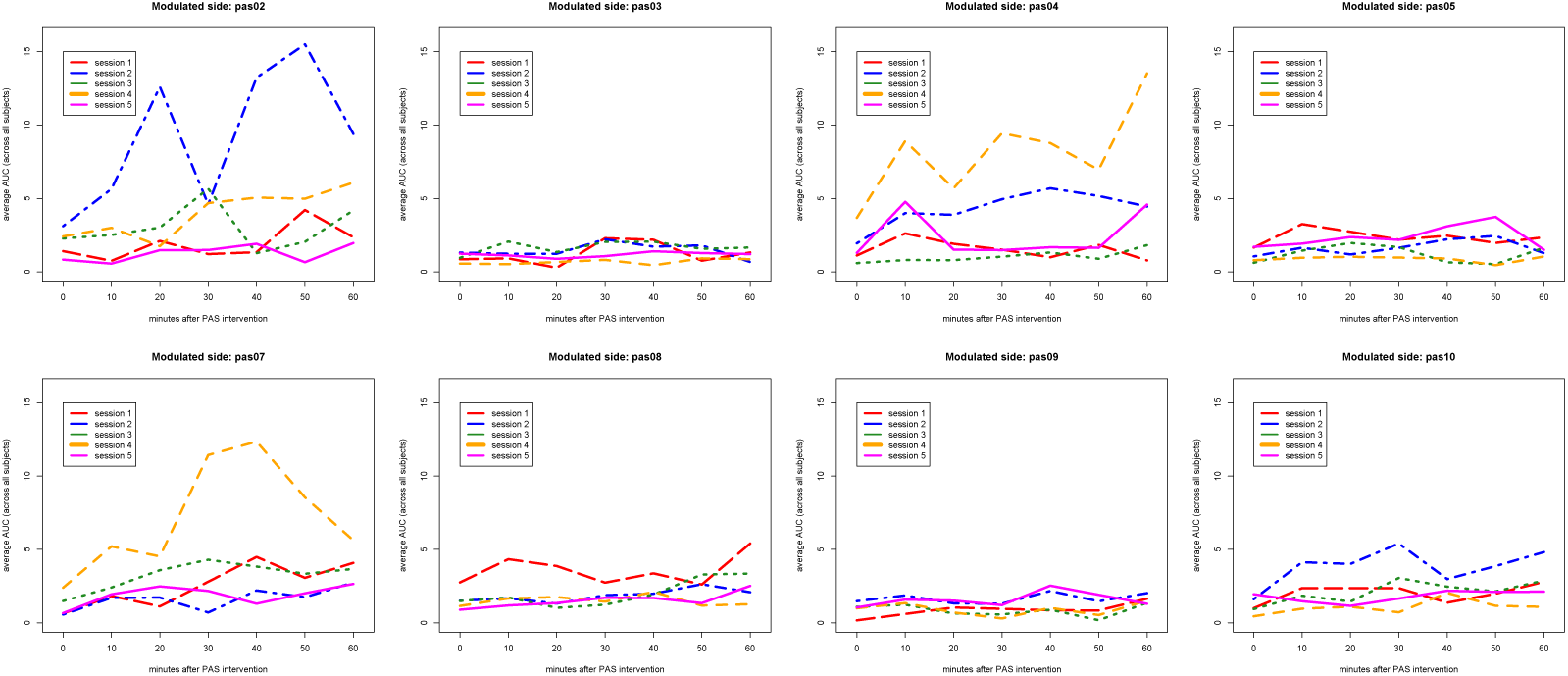
Interaction plots of each subject, modulated side only. On the individual level, all subjects showed significant time point by session interaction effects in the modulated hemisphere.

**Figure 5:**
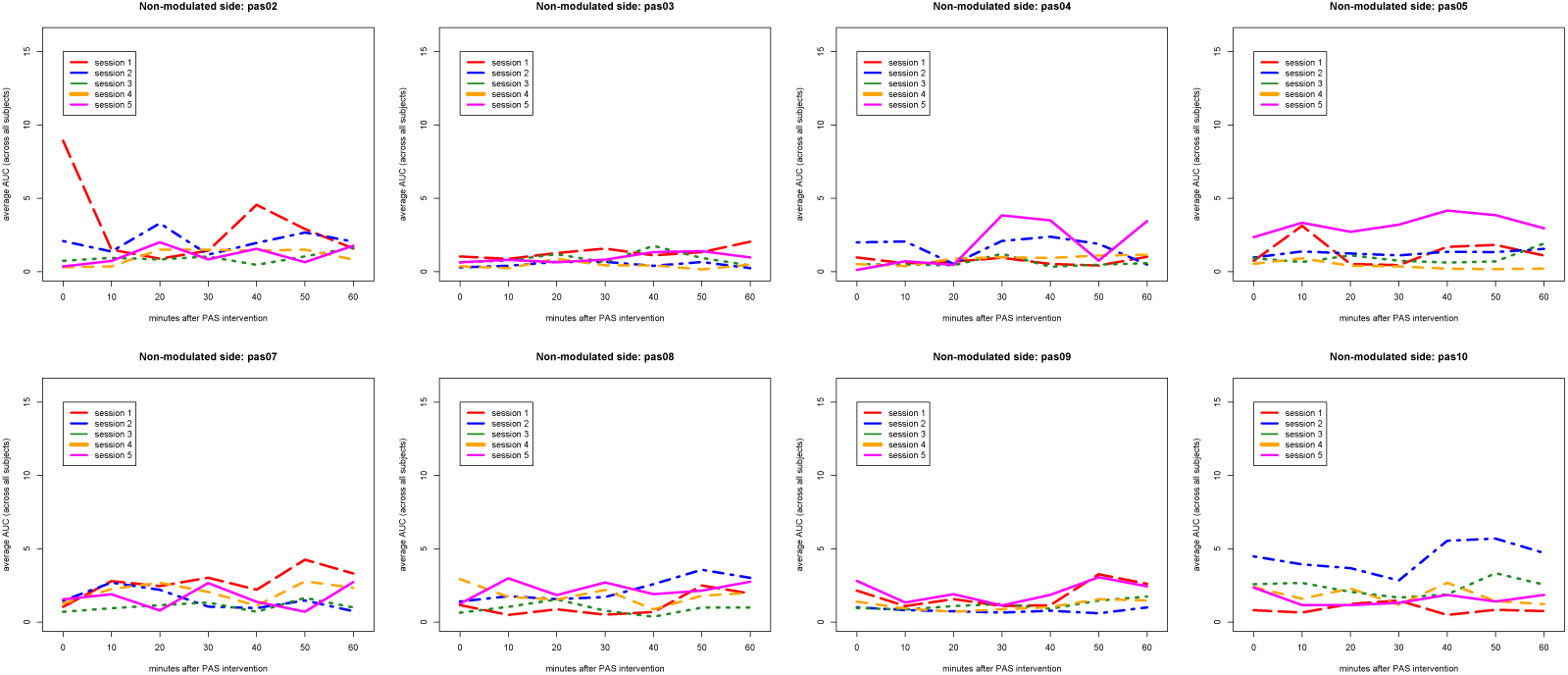
Interaction plots of each subject, non-modulated side only. On the individual level, all subjects but pas03 showed significant time point by session interaction effects in the non-modulated hemisphere.

#### 3.2.2. KS test and permutation test

For 6 out of 8 subjects, the MEPs acquired at post-PAS time points reached stability with 40 to 50 MEP measures (Figure 6). We define these subjects ‘responders’ to PAS.

**Figure 6:**
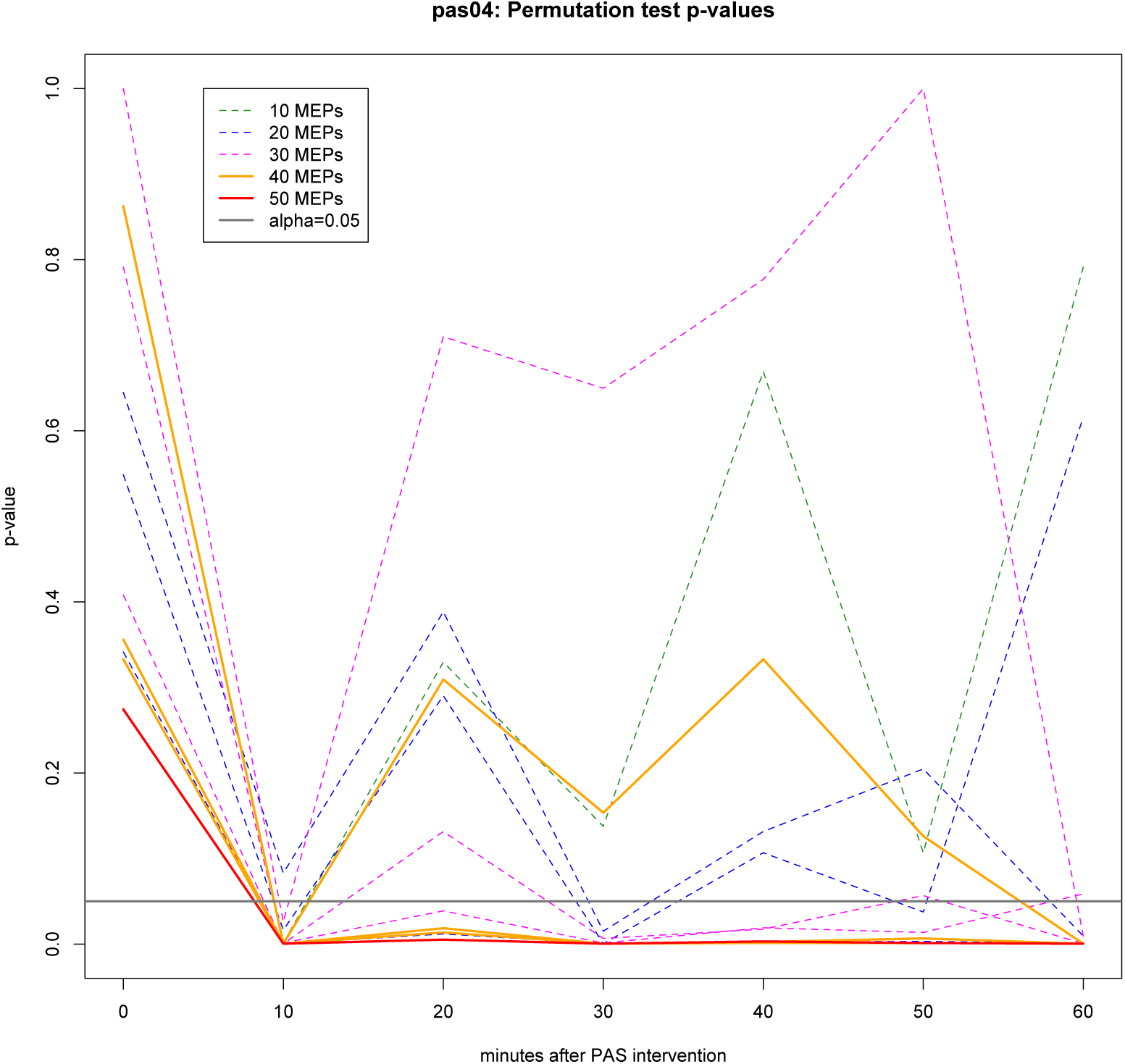
P-values (corrected) from permutation tests within subject pas04. This graph shows results from the modulated hemisphere (LM1-RFDI). At each time point after the PAS intervention, the mean differences between the post-PAS MEP data and baseline MEP data were analyzed using permutation tests. The 10 MEPs used for permutation test were selected from pas04 session 1. For 20 MEPs to 40 MEPs, three different combinations of 10 MEPs from sessions 1-5 for pas04 were studied. The results from each of these combinations of data are represented in this graph. As more MEPs are included in the permutation tests, the p-values eventually reach a level of significant change compared to baseline 10 minutes after PAS intervention.

The KS test at each post-PAS time point showed three phenomena. Three subjects (pas02, pas04, pas05) exhibited significantly increased MEP values compared to baseline at 5 or more of the 7 post-PAS time points in only the modulated hemisphere. Three subjects (pas07, pas08, pas10) showed significantly increased MEP values at 5 or more time points in both hemispheres. The remaining two subjects (pas03, pas09) displayed significant differences at only 3 or fewer time points in either hemisphere (Figures 7 and 8) and we define them ‘non-responders’ to PAS.

**Figure 7:**
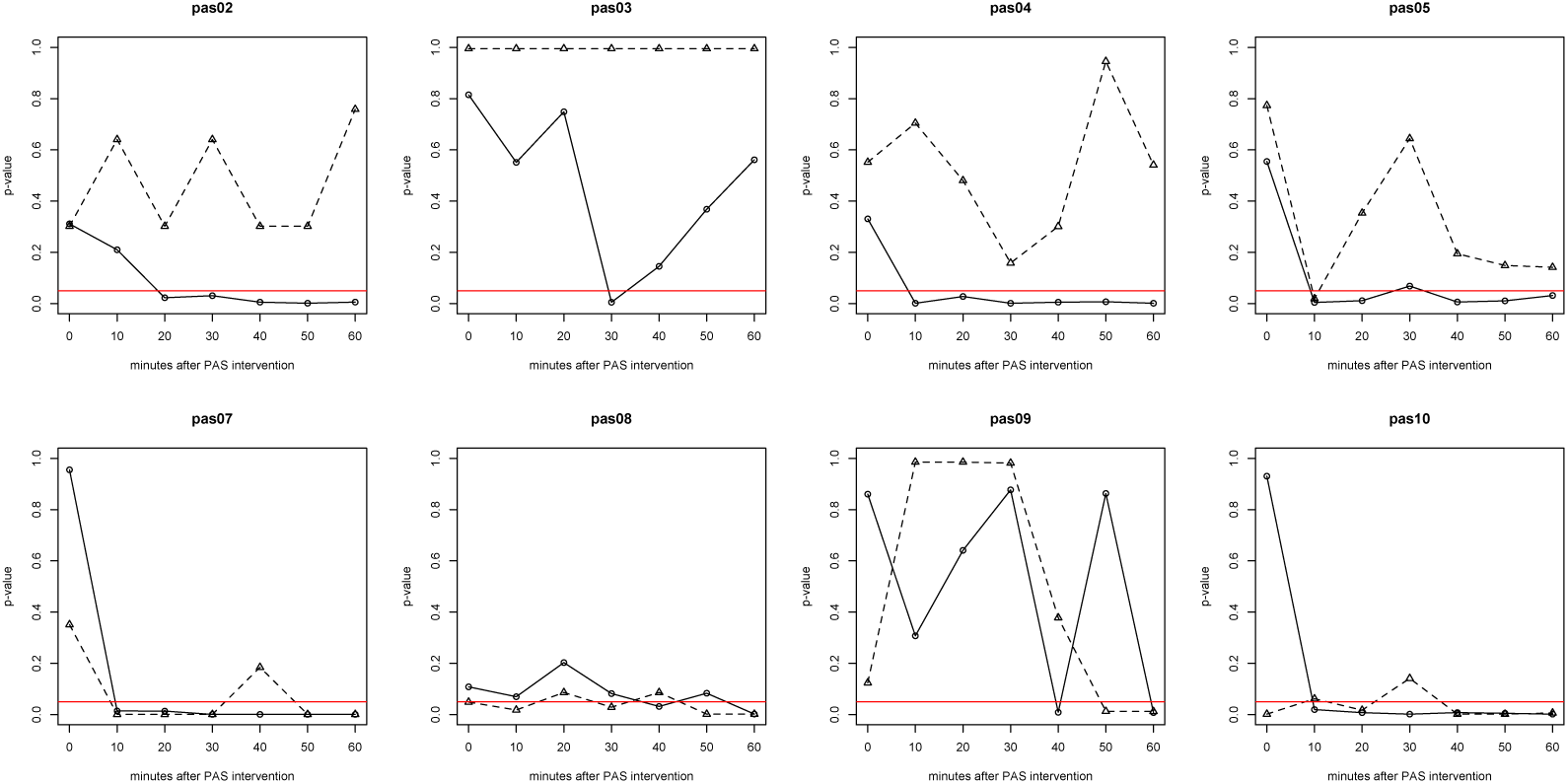
P-values (corrected) obtained from permutation tests. Data analysis included MEPs from all 5 sessions. The solid line is LM1-RFDI and the dotted line is RM1-LFDI. The red line represents alpha=0.05.

**Figure 8:**
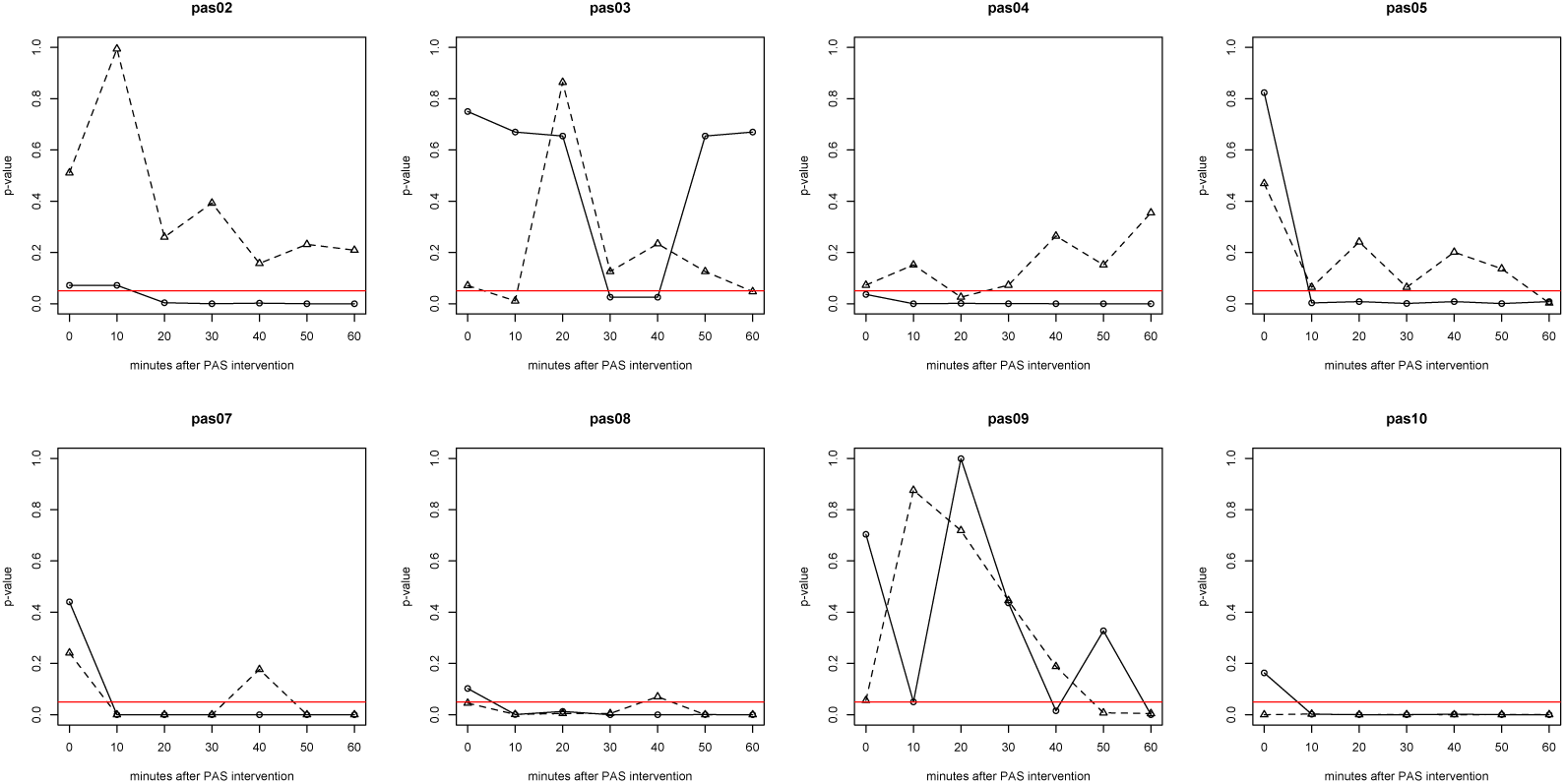
P-values (corrected) obtained from KS tests. Data analysis included MEPs from all 5 sessions. P-values from all 5 sessions. The solid line is LM1-RFDI and the dotted line is RM1-LFDI. The red line represents alpha=0.05.

As an exploratory investigation, we looked at the resting motor threshold (rMT) values in responders and non-responders. Using a permutation test, there was a significant difference in the rMT MSO% (p=0.00019996) between the 2 non-responder subjects and 6 responder subjects, with the latter cohort exhibiting a lower average rMT.

#### 3.2.3. Power factor between 40 MEPs and 50 MEPs

Since for the 6 responders out of the 8 participants, the MEPs acquired at post-PAS time points reached stability with 40 to 50 MEPs (as shown in Figure 6 for subject pas04), we compared power for 40 versus 50 MEPs. Within the modulated side, the responders (subjects having significantly increased MEPs in five or more post-PAS time points) exhibited similar power values between the 40 and 50 MEP sample sizes, with sufficient power over multiple time points except the very early ones after PAS. The non-responders (or the non-responding hemisphere in responders) demonstrated only transient increases in power at one or two time points. Power values are illustrated in figures 9 and 10.

**Figure 9:**
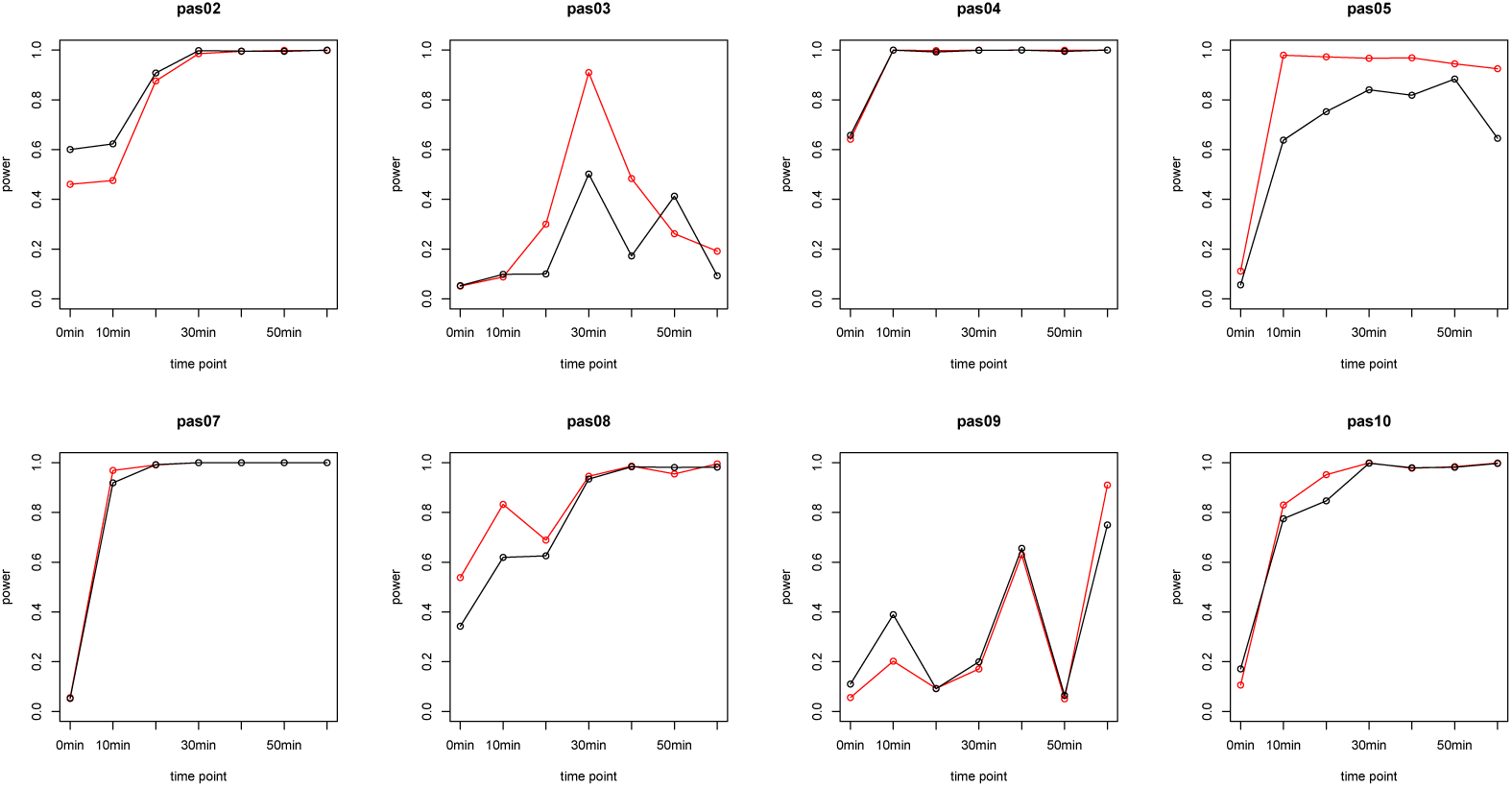
Power values from using 40 MEPs vs 50 MEPs for LM1-RFDI. The red line represents 50 MEPs, and the black line represents 40 MEPs.

**Figure 10:**
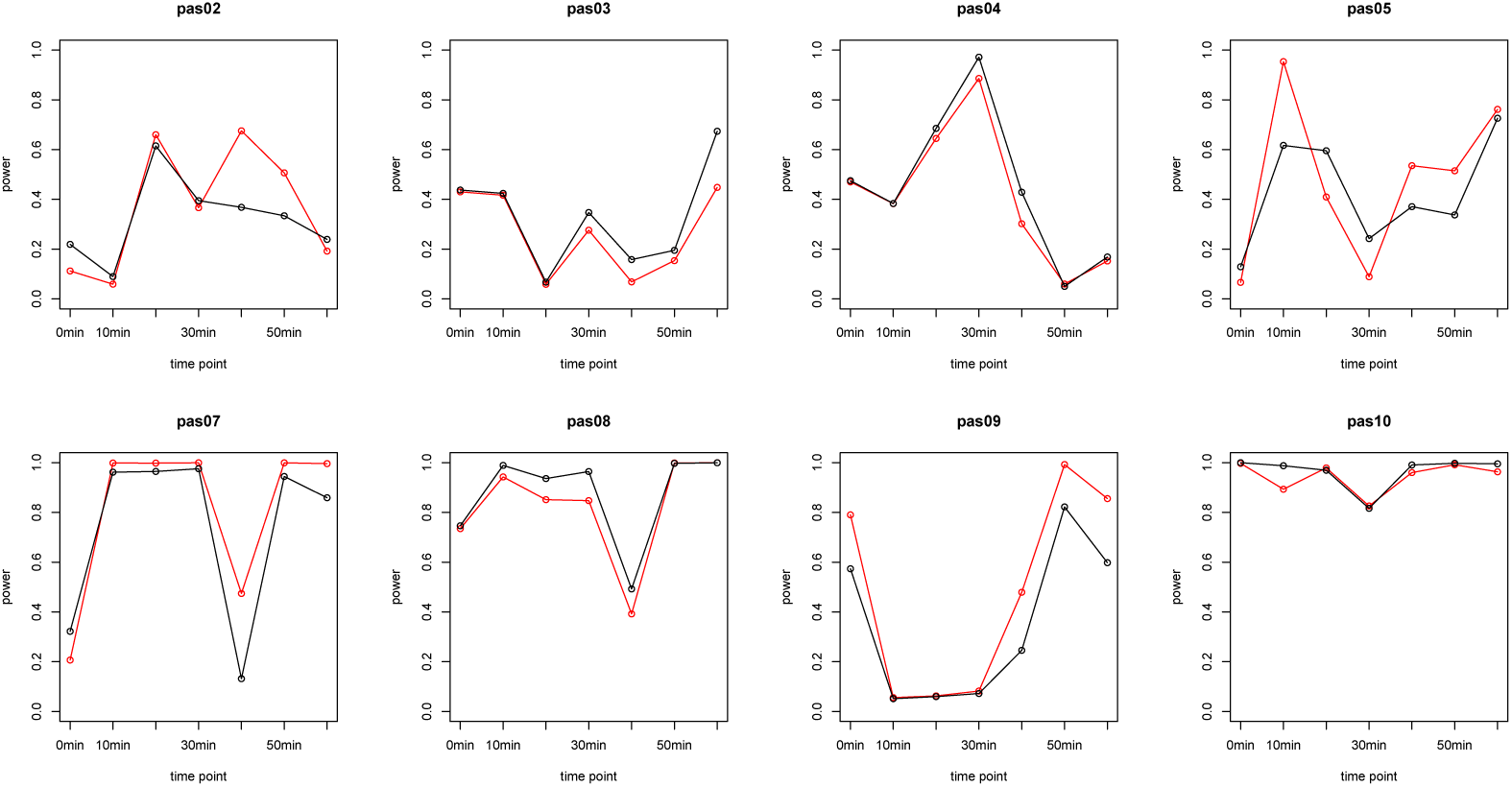
Power values from using 40 MEPs vs 50 MEPs for RM1-LFDI. The red line represents 50 MEPs, and the black line represents 40 MEPs.

A t-test and permutation test between the power measures of 40 and 50 MEPs showed that the power between these two varying numbers of samples did not statistically differ, with an exception of one time point (40 minutes post-PAS) within the modulated hemisphere. After FDR correction, only the t-test exhibited significant difference between powers of 40 MEPs and 50 MEPs at 40 minutes post-PAS. The permutation test’s p-values did not reflect this result after FDR correction (see Tables 2 and 3).

**Table 2:**
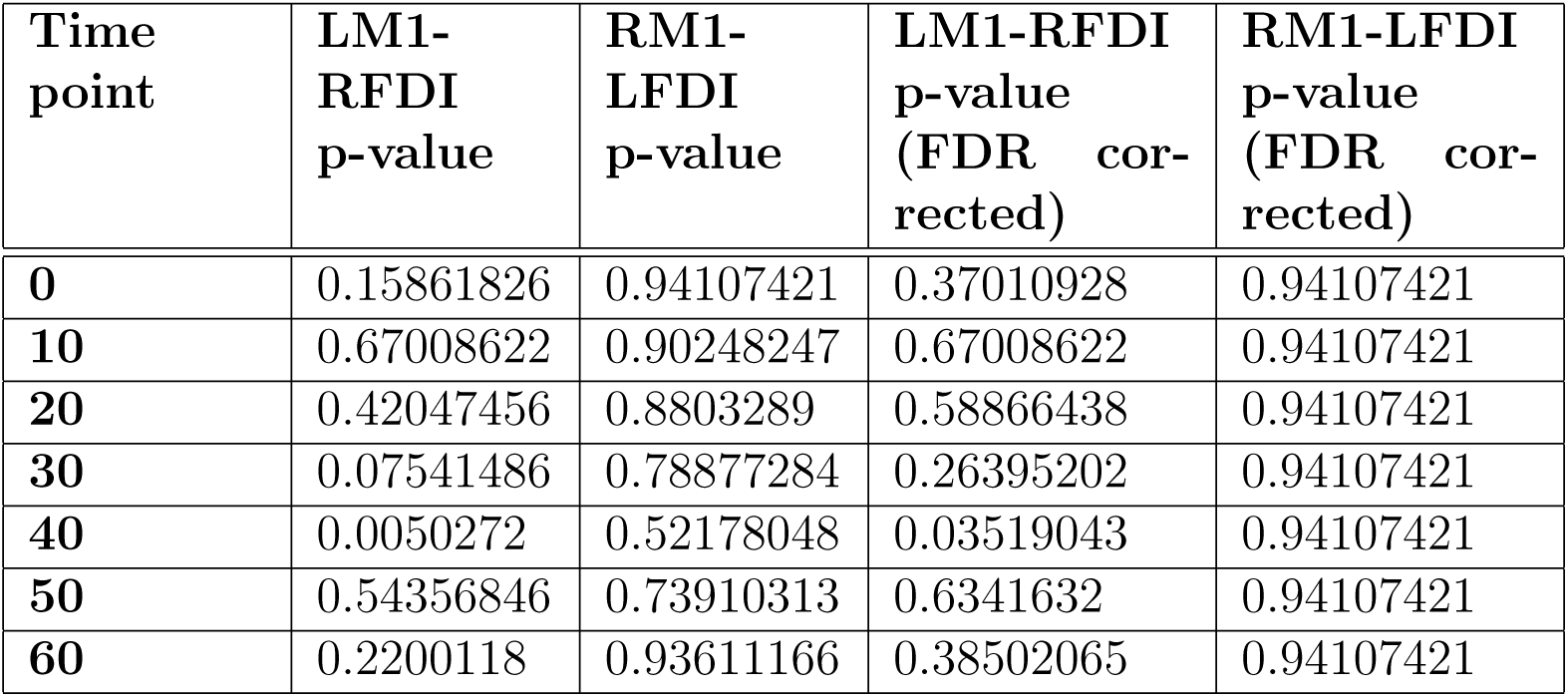
P-values acquired from t-tests testing the difference between the power values of 40 MEPs and 50 MEPs.

**Table 3:** P-values acquired from permutation tests testing the difference between the power values of 40 MEPs and 50 MEPs.

| **Time point** | **LM1-RFDI p-value** | **RM1-LFDI p-value** | **LM1-RFDI p-value (FDR corrected)** | **RM1-LFDI p-value (FDR corrected)** |
| --- | --- | --- | --- | --- |
| **0** | 0.15916817 | 0.940012 | 0.37139239 | 0.95020996 |
| **10** | 0.80123975 | 0.95020996 | 0.80123975 | 0.95020996 |
| **20** | 0.66166767 | 0.89262148 | 0.77194561 | 0.95020996 |
| **30** | 0.13357329 | 0.74885023 | 0.37139239 | 0.95020996 |
| **40** | 0.03059388 | 0.52489502 | 0.21415717 | 0.95020996 |
| **50** | 0.52009598 | 0.71045791 | 0.72813437 | 0.95020996 |
| **60** | 0.42311538 | 0.940012 | 0.72813437 | 0.95020996 |

## 4. Discussion

Due to its minimal risks and non-invasive nature, TMS is commonly employed to study neural plasticity in human subjects (Rossi et al., 2009). Paired associative stimulation, a non-invasive stimulation paradigm that comprises slow rate repetitive, paired peripheral nerve stimulation and TMS, is thought to induce LTP- or LTD-like effects. Although many studies have performed group analyses, a comprehensive assessment of the effects of PAS at individual subject level is limited. This is important since inter-individual response variability is poorly understood and confounds studies in healthy and diseased populations. Two previous studies examined intra-subject reproducibility and concluded that PAS is not reliable at individual subject level (Fratello et al., 2006; Lahr et al., 2016). However, to our knowledge, no previous study collected the amount of MEP data per subject collected and analyzed in the present study. Our results do suggest that it is possible to observe reliable individual subject level responses to PAS, provided a sufficient number of trials are collected. The results also show that these reliable individual responses do reveal individual differences in plasticity, a type of information that may be crucial for intervention studies.

Our group data from all eight subjects demonstrated a significant increase in MEP magnitude after PAS intervention only in the modulated hemisphere. This finding agrees with those from previous PAS studies, in which group effects also show PAS-induced increased cortical excitability. In addition, our group analysis exhibited no interaction between time point and session, indicating that the group response to PAS does not significantly change across sessions. This finding supports cross-sectional within-subjects PAS studies, since the PAS effect seems stable across sessions, and therefore not confounded by potential homeostatic changes with repeated exposure. This may be due to the increased signal to noise ratio from the large number of data points collected when all subjects are grouped together. Moreover, there were significant interactions between hemisphere and time point, and between hemisphere and linear trend, suggesting that the post-PAS increase in MEPs is different between the two hemispheres over time. The modulated hemisphere exhibited a strong linear trend in MEP increases over time, while the non-modulated hemisphere only approached significance.

In contrast, at an individual level, we observed significant interactions between time point and session. Since the post-PAS MEPs were normalized, these interactive effects suggest that the post-PAS MEPs exhibited different temporal variations between sessions. This variability can be attributed to both physiological and/or technical factors.

Previous studies have found a number of physiological factors that may be associated with the variability of PAS-induced plasticity response within an individual. For example, a subject’s attentional state during the intervention has been reported to significantly influence MEP magnitude. Stefan et al. (2004) found that a greater amount of plasticity was induced when the participants focused on their target hands versus on a diverting cognitive task during the intervention. This may be due to the deactivation of noncognitive-related areas of the brain during cognitive tasks, possibly affecting the efficacy of the neuroplasticity processes (Stefan et al., 2004; Conte et al., 2007, 2008; Antal et al., 2007). Although our protocol asks participants to focus on the target muscle, their attention may have varied from session to session. Moreover, the time of day may affect neural plasticity (Sale et al., 2007, 2008; Marshall et al., 2004). In a multiple sessions study like this one scheduling conflicts make it difficult to study subjects always at the same time of day.

Multiple technical elements could also contribute to the individual variability of MEP response. First, the location, approach angle, and rotational angle of the coil significantly influence the MEP response (Brasil-Neto et al., 1992; Richter et al., 2013; Kiers et al., 1993). Brasil-Neto et al. (1992) demonstrated that the optimal direction of the magnetically induced currents is orthogonal to the central sulcus, while Silva et al. (2008) hypothesized that because the current is parallel to the cortical surface, the interneurons lying perpendicular to the central sulcus are preferentially activated. In addition, Kiers et al. (1993) found that the variability of MEP response was inversely correlated with stimulus intensity and the recruitment of motor neurons, amongst other factors. The protocol used here, as in many other studies, required to manually reposition the TMS coil at each time point, perhaps introducing minor deviations in coil positioning that may have contributed further to the inconsistency of MEP measures across the individual time points of the sessions. In an effort to reduce some of these technical issues, we used BrainSight TMS Neuronavigation at each session to obtain a visual feedback of the TMS coil location in respect to BrainSight’s digitized spatial coordinate system, increasing targeting precision and reliability, and effectiveness of neuromodulation (Bashir et al., 2011).

Considering that PAS-induced MEPs changed significantly across sessions, we chose to investigate the number of MEPs necessary to reach a level of reliability within one subject. To mimic the increase in the signal to noise ratio seen in group data (and likely due to a higher number of data points in group analyses), we chose a statistical approach for individual analyses in a similar fashion. Using varying combinations of post-PAS MEP measures, we applied permutation and KS tests at each post-PAS time point against the baseline MEP measures. As shown in Figure 6, we observed that the behavior of the subjects’ MEPs became more reliable with 40 to 50 MEPs per time point. With this method, the participants displayed three patterns of response: (1) those who responded in both hemispheres, (2) those who responded only in the modulated hemisphere, (3) and those who did not respond, wherein responsiveness was defined as a significant deviation from baseline values.

This result is a step forward in reliably assessing PAS-induced neural plasticity at individual subject level. Such assessment may have valuable applications in the clinical domain. However, it is not very practical to have an assessment that requires multiple visits. Thus, future studies should assess whether our results are replicable in a single session that collects at least between 40 to 50 MEPs per time point to reliably establish a plasticity response at the individual subject level.

As mentioned previously, age can potentially contribute to individual variability, though results so far are mixed: whereas several studies have shown that cortico-spinal excitability decreases with age, trial-to-trial variability of MEPs has been reported to increase with age (Müller-Dahlhaus et al., 2008; Tecchio et al., 2008; Fathi et al., 2010; Todd et al., 2010; Pitcher et al., 2003). While our cohort was composed of young adults with a fairly narrow range (range = 19-26, M = 24.4, SD = 6.15), we contend that the main difference between the current study and previous ones is in the number of MEPs collected in each subject. To better assess age effects on individual differences and reliability, studies should collect the number of MEPs per subject per time point that our analyses suggest.

An exact binomial test of the distribution of gender in our sample (2 males, 6 females) exhibited a p-value of 0.2841. Due to our small sample size, we chose not to evaluate significant differences according to gender. Visually, we did not detect any differences in MEP response between our male and female subjects, but Pitcher et al. (2003) have found that females demonstrate larger MEP variability, possibly influenced by fluctuating levels of ovarian steroid hormones during their menstrual cycles (Smith et al., 1999; Wassermann, 2002).

The level of physical activity performed by an individual can also affect the plasticity in the motor cortex. Cirillo et al. (2009) have shown that the motor cortex plasticity, measured by the MEP amplitude, is significantly enhanced in long-term physically active individuals, who engaged in more than 150 minutes of moderate-to-vigorous aerobic activity per day on at least five days a week, when compared to sedentary participants, who performed less than 20 minutes per day of exercise on less than 4 days per week. We did not record our participants’ frequency and intensity of their regular exercise routine. This may have affected both individual differences between responders and non-responders and the session to session variability within individuals. This does not invalidate our method for extracting reliable individual differences in response to PAS, since many factors do affect plasticity and it is highly unlikely that individual studies can control all these factors.

Our exploratory finding of a lower average rMT in responders is in agreement with a previous study, which found responders to have a significantly lower rMT (Müller-Dahlhaus et al., 2008). The relationship between rMT and response to PAS is obviously interesting and future studies with larger cohorts should investigate it further.

Recent findings suggest that genetics may also play a role in PAS-induced neuroplasticity. In Cheeran et al. (2008) and Kleim et al. (2006), participants with the brain-derived neurotrophic factor (BDNF) polymorphism Val66Met exhibited lower PAS-induced motor cortex plasticity compared to those with Val66Val. The authors hypothesize that this is due to BDNF’s effects on synaptic growth and differentiation during LTP/LTD. Other studies have suggested that TMS-induced brain plasticity is genetically inheritable (Missitzi et al., 2011; Pellicciari et al., 2009). Our study cannot tell whether the individual differences in neural plasticity we observed are linked to genetic differences, but undoubtedly having a way to reliably assess individual differences in neural plasticity is a step forward to better understand the genetic underpinnings of neural plasticity.

We also observed individual differences in PAS response to the non-modulated hemisphere. Three of our ‘responders’ do show reliable changes in the right hemisphere. The practice of probing the non-modulated hemisphere is not common in PAS studies. Very little is known about these changes even at group level. Future studies should investigate the potential factors underlying these responses in the non-modulated hemisphere. In responders, MEPs began to significantly change approximately 10 to 20 minutes after PAS modulation, supporting previous findings (Tecchio et al., 2008; Müller-Dahlhaus et al., 2008; Ziemann et al., 2004). Our data suggest that subject level PAS studies should focus on measuring post-PAS effects after 10 or 20 minutes. Since power does not seem to change significantly between 40 and 50 MEPs, 40 MEPs may be sufficient to discern individual plasticity profiles at each time point, but collecting 50 MEPs is certainly safer. With a reproducible PAS response at individual subject level, it becomes possible to examine the potential for augmentation of the PAS response by a drug or rehabilitation therapy that alters excitability or synaptic efficacy at the onset of a trial. Thus, PAS may serve as a predictive biomarker of the intervention’s potential effect on adaptive plasticity for the upper extremity after, for example, hemiparetic stroke (Kim and Winstein, 2017; Cramer et al., 2011). It may also be feasible to monitor changes in the PAS response over the time of therapy to better understand whether the motor hand region is being engaged or affected by the intervention in the direction of greater plasticity and behavioral improvement.

## 5. Conclusion

Our study shows that PAS may be a reliable tool in assessing individual neural plasticity responses, provided a sufficient number of data points is collected. Information on individual propensity to neural plasticity would be desirable as a biomarker when testing targeted, individualized therapies for motor impairments. This study provides an important step toward achieving that goal. However, practical considerations make the multi-session approach of this study a burden on participants. Future studies should assess whether our findings can be replicated with sufficient data collected in one session only, a more desirable approach for innovative clinical research.

